# Isolation and quantification of bacterial membrane vesicles for quantitative metabolic studies using mammalian cell cultures

**DOI:** 10.1101/2023.09.25.559254

**Authors:** Marcel Kretschmer, Julia Müller, Petra Henke, Viktoria Otto, Alejandro Arce Rodriguez, Mathias Müsken, Dieter Jahn, José Manuel Borrero-de Acuña, Meina Neumann-Schaal, Andre Wegner

**Affiliations:** Department of Bioinformatics and Biochemistry, Technische Universität Braunschweig, Rebenring 56, 38106 Braunschweig, Germany; Leibniz Institute DSMZ - German Collection of Microorganisms and Cell Cultures, Inhoffenstraße 7 B, 38124 Braunschweig, Germany; Institute for Microbiology, Technische Universität Braunschweig, Braunschweig, Germany; Braunschweig Integrated Center of Systems Biology (BRICS), Technische Universität Braunschweig, Rebenring 56, 38106 Braunschweig, Germany; Central Facility for Microscopy, Helmholtz Centre for Infection Research (HZI), Inhoffenstraße 7, 38124 Braunschweig, Germany; Department of Microbiology, Facultad de Biología, University of Sevilla, Av. de la Reina Mercedes 6, Sevilla, CP 41012, Spain

**Author notes:** These authors contributed equally. **Corresponding Author** Andre Wegner.

**Keywords:** bacterial membrane vesicles (BMVs), membrane vesicles (MVs), outer membrane vesicles (OMVs), vesicle isolation, quantification, *Pseudomonas aeruginosa*, pathogen, SuhB, metabolism

## Abstract

Bacterial membrane vesicles (BMVs) are produced by most bacteria and participate in various cellular processes, such as intercellular communication, nutrient exchange, and pathogenesis. Notably, these vesicles can contain virulence factors, including toxic proteins, DNA, and RNA. Such factors can contribute to the harmful effects of bacterial pathogens on host cells and tissues. Although the general effects of BMVs on host cellular physiology are well known, the underlying molecular mechanisms are less understood. In this study, we introduce a vesicle quantification method, leveraging the membrane dye FM4-64. We utilize a linear regression model to analyze the fluorescence emitted by stained vesicle membranes to ensure consistent and reproducible vesicle-host interaction studies using cultured cells. This method is particularly valuable for identifying host cellular processes impacted by vesicles and their specific cargo. Moreover, it outcompetes clearly unreliable protein concentration-based methods. We (1) show a linear correlation between the quantity of vesicles and the fluorescence signal emitted from the FM4-64 dye, (2) introduce the “vesicle load” as a new semi-quantitative unit, facilitating more reproducible vesicle-cell culture interaction experiments (3) show that a stable vesicle load yields consistent host responses when studying vesicles from *Pseudomonas aeruginosa* mutants (4) demonstrate that typical vesicle isolation contaminants, such as flagella, do not significantly skew the metabolic response of lung epithelial cells to *P. aeruginosa* vesicles, and (5) identify inositol-1-monophosphatase (SuhB) as a pivotal regulator in the vesicle-mediated pathogenesis of *P. aeruginosa*.

## INTRODUCTION

*Pseudomonas aeruginosa* is an opportunistic pathogen that typically targets the respiratory tract of immunocompromised individuals, causing infections which in their chronic state can ultimately lead to fatal lung disease, in particular in patients with cystic fibrosis [1, 2]. In the context of *P. aeruginosa* infections, the release of bacterial membrane vesicles (BMVs) has been recognized to contribute to bacterial pathogenesis. BMVs provide the means to deliver a diverse bacterial cargo into host cells, including nucleic acids, proteins, and metabolites [3]. While the general effects of BMVs on cellular physiology have been studied in detail [4–10], the underlying molecular mechanisms are less understood. However, understanding these mechanisms is critical to pinpoint new drug targets or develop new treatment strategies for *P. aeruginosa* infections. Methods to analyze vesicle-related disease mechanisms often rely on quantitative readouts of the infected host cell, including changes in gene expression, cell proliferation, or cellular metabolism. These methods are highly sensitive to the vesicle amount and corresponding bacterial cargo which is applied to the host cells. In this context, exact vesicle quantification is crucial for the correct interpretation of data obtained from vesicle-host interaction studies. However, multiple factors such as culture conditions, the bacterial strain, and especially the used BMV isolation procedure lead to significant differences in the amount and composition of vesicles [11]. Therefore, it is important to determine the precise vesicle amount to perform reproducible mammalian cell culture experiments with isolated bacterial vesicles. Several quantification methods have been developed in the past, either based on vesicle cargo or quantity [11]. Among these, quantification by protein concentration is widely used, as it is a relatively simple and established method in most laboratories [11]. However, the protein concentration might vary independent of the vesicle number or size. For example, non-vesicle proteins (e.g. flagella or proteins from lysed bacteria) are the most common contaminants during vesicle isolation, making vesicle-host interaction experiments with normalized protein concentrations often not reproducible [12]. A more suitable approach for BMV quantification is nanoparticle tracking analysis which determines the vesicle number and size distribution [12]. However, most vesicle-host interaction experiments require a quantification protocol that delivers an equal amount of pathogenic cargo to the host cell, e.g. for the identification of specific vesicle factors important for the infection (e.g., proteins, RNA, DNA, or metabolites). Since nanoparticle tracking analysis is normalized by the vesicle number, it can provide a different cargo amount depending on the vesicle size. A third method for vesicle quantification is the staining of vesicle membranes with a lipid dye (e.g. FM4-64) and subsequent detection of the emitted fluorescence [11, 13, 14]. This quantification method has the advantage that it more robustly delivers an equal amount of pathogenic cargo, because the emitted fluorescence will depend on both, the number and size of vesicles, but is independent of specific cargo (e.g. protein concentration). Nevertheless, FM4-64 has been employed solely to compare the relative quantity of vesicles between distinct bacterial strains and has not been utilized to modulate the vesicle amount in cell culture interaction experiments.

Here, we present a vesicle quantification method suitable to reproducibly analyze the effect of bacterial vesicles on cellular host processes. The method is based on the membrane dye FM4-64 and a linear regression model of the emitted fluorescence. We show that this method is particularly well-suited to analyze the vesicle-induced effect on host cell metabolism.

## METHODS

### Bacterial strains

The *P. aeruginosa* PA14 wild-type strain [15] and several PA14 MAR2xT7 transposon library insertion mutants [16] were used throughout this study. In addition a PA14 *fliC* knockout mutant strain missing functional flagella was used [17].

### Cultivation of bacterial strains

*P. aeruginosa* PA14 [15], which shows an increased virulence potential compared to the commonly used strain PAO1 [18], and it’s derived mutant strains were used in this study. For BMV isolation, strains of interest were inoculated in 20 mL lysogeny broth (LB, Roth) with additional 50 µg/mL gentamicin (Roth) for transposon mutant strains (Table 1) and incubated at 37 °C and 140 rpm overnight. The optical density (OD) of pre-cultures was measured at a wavelength of 600 nm. Main cultures were inoculated with an OD600 of 0.05 in 100 mL LB broth. To gather a sufficient amount of isolated BMVs, a final volume of at least 400 mL is recommended. Cultivation was performed with flasks with baffles and a 1:5 volume ratio (culture:flask). Main cultures were incubated at 37 °C with 160 rpm until they reached the early-stationary phase of growth. Individual growth phases of each strain were determined beforehand.

**Table 1:** Bacterial strains used in this study.

| *P. aeruginosa* strain | Features | Source |
| --- | --- | --- |
| PA14 wild-type | Parental strain | [15] |
| PA14 *glpQ* mutant | PA14 *glpQ* transposon mutant strain (MAR2 x T7) | [16] |
| PA14 *suhB* mutant | PA14 *suhB* transposon mutant strain (MAR2 x T7) | [16] |
| PA14 *lap* mutant | PA14 *lap* transposon mutant strain (MAR2 x T7) | [16] |
| PA14 *lipA* mutant | PA14 *lipA* transposon mutant strain (MAR2 x T7) | [16] |
| PA14 *oprF* mutant | PA14 *oprF* transposon mutant strain (MAR2 x T7) | [16] |
| PA14 Δ*fliC* mutant | PA14 *fliC* knockout mutant strain | [17] |

### Membrane vesicle isolation

BMVs were isolated from supernatants of bacterial cultures by ultrafiltration combined with ultracentrifugation. Main cultures were used upon reaching the early-stationary growth phase. Bacterial cells were removed by centrifugation at 4 °C and 8,000 ˗ g for 30 min and two filtrations with 0.22 µm polyethersulfone (PES) filters (Corning^TM^, 10268951). Cell-free supernatants were concentrated to 1 mL by ultrafiltration through a membrane with a molecular weight cut-off of 30 kDa (VivaSpin 20, 30,000 MWCO PES (Sartorius, VS2022)) at 4 °C and 4,000 ˗ g. The remaining material was resuspended in 6 mL sterile 1× PBS (Gibco). Samples were ultracentrifuged at 4 °C and 150,000 ˗ g for 2 h (Optima L-80 XP Ultracentrifuge with Type 70.1 Ti Fixed-Angle Titanium Rotor (Beckman Coulter)) and supernatants were removed. The remaining vesicle pellets were resuspended in 2 mL sterile 1× PBS and sterile filtered with 0.20 µm syringe filters (Carl Roth, Sartorius VS0271).

### Membrane vesicle purification with density gradient centrifugation

Bacterial main cultures (10x 200 mL) were inoculated with an OD600 of 0.05 and harvested in their early-stationary phase. Bacterial cells were removed by centrifugation at 4 °C and 8,000 ˗ g for 30 min and sterile filtered through 0.22 µm PES filters (Corning^TM^, 10268951). The cell-free supernatant (CFS) was concentrated to 10 mL with a 100 kDa ultrafiltration cassette (Sartorius, VF05H4) attached to a peristaltic pump, followed by another centrifugation at 8,000 ˗ g for 30 min and 0.22 µm sterile filtration. The concentrated CFS was then ultracentrifuged at 100,000 ˗ g and 4 °C for 3 h (Optima L-80 XP Ultracentrifuge with Type 70.1 Ti Fixed-Angle Titanium Rotor (Beckman Coulter)). The pellet was resuspended in 1 mL of 20 mM HEPES / 100 mM NaCl, pH 7. An iodixanol (OptiPrep, Sigma-Aldrich) density gradient was used to remove remaining soluble proteins and impurities. Samples were diluted to 50 % (v/v) OptiPrep-HEPES/NaCl and 2 mL were transferred to the bottom of the tube. The latter was then sequentially layered with 2 mL of 40 %, 2 mL of 35 %, 4 mL of 30 % and 2 mL of 25 % OptiPrep-HEPES/NaCl (adapted, [19]). The gradient was centrifuged at 4 °C and 100,000 ˗ g for 18 h (Optima L-80 XP Ultracentrifuge with Swing Rotor (Beckman Coulter)). After centrifugation 500 µL fractions were transferred from top to bottom into a new reaction tube and enumerated accordingly. The fractions 13 and 20 were used representatively for experiments as “pure” and “impure” fractions respectively. Lipid and protein concentrations were measured using FM4-64 lipid dye (Invitrogen) and BCA protein assay (Thermo Scientific, 23225).

### Vesicle and protein quantification

Yields of membrane vesicles were determined by the fluorescence signal emitted by the membrane lipid dye FM4-64 (Invitrogen, T13320). Stock solutions were prepared in water at a concentration of 50 µg/mL and kept at 4 °C. Before every measurement a working solution for staining was prepared and kept on ice: The stock solution was diluted to a final concentration of 5 µg/mL in Hanks Balanced Salt Solution (HBSS) containing 0.4 g/L KCl, 0.06 g/L KH2PO4, 8 g/L NaCl, 0.05 g/L Na2HPO4 and 1 g/L glucose. The staining was performed with a 100-fold dilution of the sample in the working solution. For each measurement, 200 µL were then transferred into one well of a 96-well plate (Greiner, transparent, flat bottom). The fluorescence was measured immediately with a plate reader (Tecan Spark) using the following settings: excitation at 558 nm, emission at 734 nm, gain = 150, 30 flashes, Z-position = 20000, and room temperature. Data were obtained from 3 technical replicates.

Protein concentrations of membrane vesicle samples were determined by using a BCA protein assay (Thermo Scientific, 23225). For the microplate procedure, 25 µL of each sample were used as described in the user guide. Data were obtained from 2 technical replicates.

### Cell confluence

Confluence measurements were performed with a plate reader (Tecan Spark) at 37 °C. Cells were pre-incubated at standard culture conditions in Dulbecco’s Modified Eagle Medium (DMEM, Gibco, 41965039) containing additional 10 % dFBS (bio-sell, FCS.DIA.0.500) for 24 h before the first confluence measurement (0 h). Right before treatment, cells were washed with 1× PBS. Vesicles or flagellin were diluted in standard culture medium, followed by a three-day incubation before the second confluence measurement (72 h). The relative confluence change was determined on a per well basis by dividing with the starting confluence (0 h).

### Stable isotope labeling

Cells were seeded with 200,000 cells per well in 6-well plates (Greiner, transparent, flat bottom) and pre-incubated at standard culture conditions for 24 h. The medium was then discarded and cells were washed with 1× PBS. 10 % dFBS (bio-sell, FCS.DIA.0.500), 25 mM [U-^13^C_6_]glucose (Cambridge Isotope Laboratories, Inc., CLM-1396-1), and 4 mM glutamine were added to SILAC DMEM medium (Gibco, A14430-01). New medium and *P. aeruginosa* BMVs were applied to cells and incubated for 24 h. Cell confluence was measured before metabolite extraction.

### Metabolite extraction and GC/MS analysis

Cells were washed with 0.9 % NaCl and quenched with 400 µL ice-cold methanol and 400 µL ice-cold water containing 1 µg/mL [6-D_1_]glutaric acid (internal standard). Cells were detached from the well surface with a cell scraper. The cell suspension was then transferred into a tube with 400 µL ice-cold chloroform and mixed at 1,400 rpm and 4 °C for 20 min, followed by centrifugation at 17,000 ˗ g and 4 °C for 5 min. For analysis of polar metabolites, 200 µL of the aqueous phase of each sample were transferred into a glass vial with an inlet and dried in a vacuum centrifuge at 4 °C. The vials were air-tight sealed. For targeted metabolite analysis with GC-MS, each dried polar metabolites sample was dissolved in 15 µL of 2 % methoxyamine in pyridine and mixed at 55 °C for 90 min. After dissolution, *tert* -butyldimethylsilyl (TBDMS) derivatization was done with 15 µL of *N*-tert-butyldimethylsilyl-*N*-methyltrifluoroacetamide (MTBSTFA) for another 60 min at 55 °C under continuous shaking. GC-MS analysis was performed with an Agilent 7890A GC equipped with a 30 m ZB-35 capillary column (Phenomenex) and a 5 m guard capillary column coupled to an Agilent 5975C inert mass selective detector (Agilent Technologies). Electron impact ionization (EI) was carried out at 70 eV. One microliter of each sample was injected into the SSL injector in splitless mode at 270 °C operating with helium as a gas carrier (total flow = 1 mL/min). GC oven temperature was heated to 100 °C for 2 min and increased to 300 °C at 10 °C/min. After 4 min the oven was heated to 330 °C. The temperature at the MS source and quadrupoles was held at 230 and 150 °C, respectively. The detector was run in selected ion monitoring (SIM) mode. Mass isotopomer distributions were determined by integration and corrected for natural isotope abundance with the data analysis software Metabolite Detector (https://md.tu-bs.de).

### Electron Microscopy

A thin carbon film was floated on 30 - 50 µl sample droplets to allow adhesion of vesicles. After one minute, a 300 mesh copper grid was used to lift off the carbon film. The grid was washed twice on droplets of distilled water and placed on a droplet of 4 % uranylacetate (w/v). After one minute, the excessive liquids were carefully removed with a filter paper and the grids were heat-dried with a 60 W light bulb. Samples were examined with a Libra 120 transmission electron microscope (Zeiss, Germany) with an acceleration voltage of 120 kV and at calibrated magnifications. Contrast and brightness adjustments as well as size measurements were done with the WinTEM software.

## RESULTS AND DISCUSSION

This study aimed at the identification of host cell processes influenced by *P. aeruginosa* vesicles as well as the identification of the responsible factors within the vesicles (Figure 1). One sound strategy to e.g. identify important pathogenic vesicle proteins is the use of specific mutants that lack abundant vesicle proteins and compare host responses to that induced by wild-type vesicles. In case that an important pathogenic protein is missing, one would expect a changed host response. For example, we used a *P. aeruginosa* glycerophosphoryl diester phosphodiesterase gene transposon mutant (*glpQ*) which inactivates the gene encoding for one of the most abundant proteins in *P. aeruginosa* vesicles. As BMVs derived from different mutant strains may differ in their composition, it is mandatory for host-vesicle interaction experiments to quantify the precise vesicle amount given to the cells independent of their specific cargo. To that end, we introduce a quantification method based on a vesicle staining with the membrane dye FM4-64, which allows us to perform reproducible vesicle-host interaction experiments with BMVs isolated from different *P. aeruginosa* strains.

**Figure 1:**
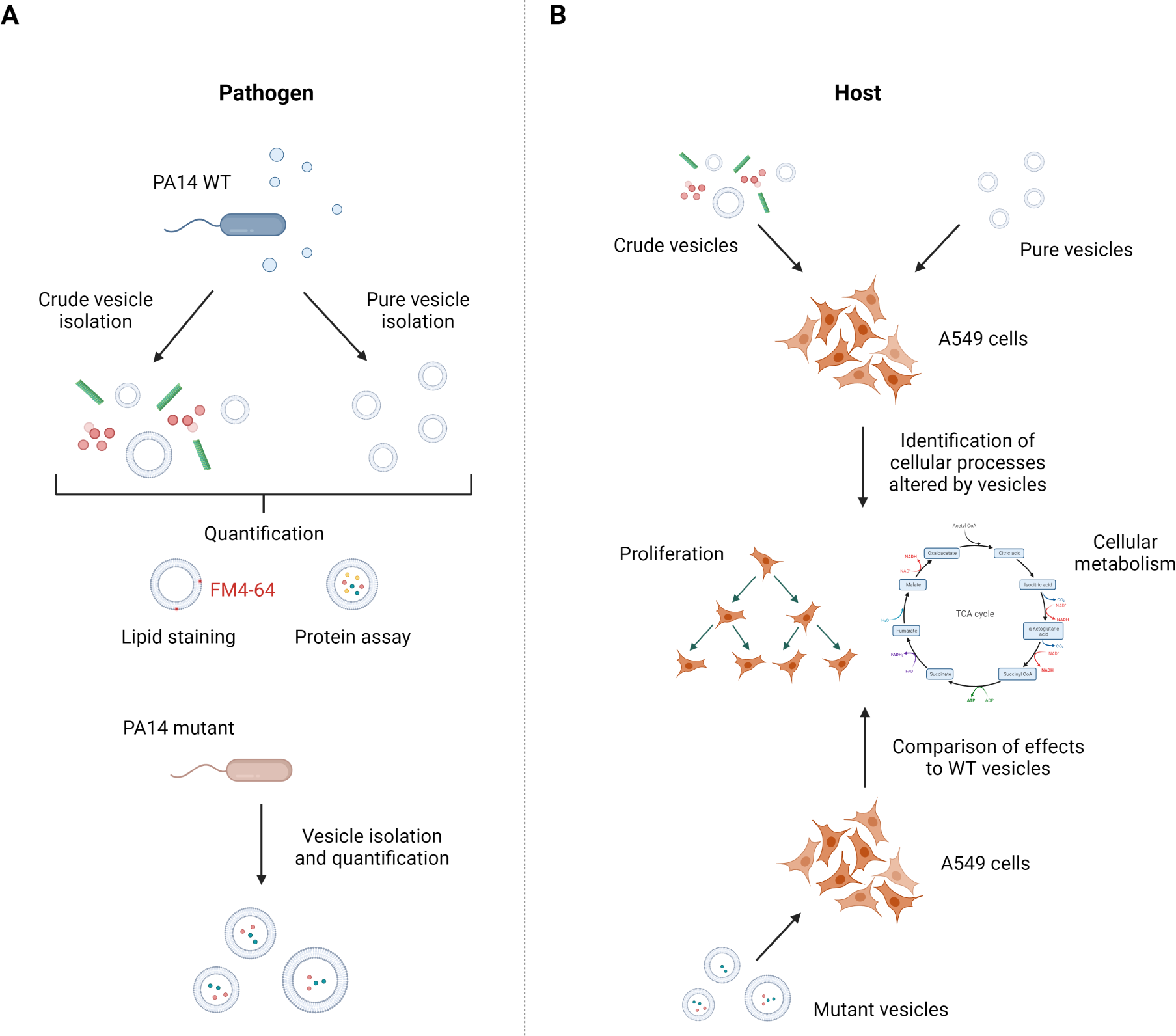
Isolation and quantification strategy of *P. aeruginosa* PA14 vesicles to identify vesicle-mediated cellular processes in A549 cells. (A) To compare the effect of different BMV pools, crude and pure vesicles from *P. aeruginosa* wild-type and several mutant strains were isolated and subsequently quantified by lipid content and protein concentration. (B) A549 lung cancer cells were treated with crude and pure vesicles from *P. aeruginosa* wild-type strain to analyze cellular processes that are affected by BMVs like proliferation and cellular metabolism. To identify responsible vesicle factors for the observed effects, A549 cells were treated with mutant vesicles missing abundant BMV proteins and changes in cellular processes compared with the effects of wild-type vesicles. Figure created with biorender.com.

To show that FM4-64 staining is suitable for producing consistent and reproducible vesicle-host interaction studies, we isolated BMVs from *P. aeruginosa* using two protocols: One relatively fast method to isolate “crude” vesicles with impurities (Figure S1A) and one based on density gradient centrifugation to isolate vesicles almost free of impurities (Figure S1B). We used transmission electron microscopy (TEM) to assess the quality and purity of all isolated vesicles (Figure S2). As expected, crude wild-type BMVs isolated with the fast method show impurities like pyocins and flagella debris (Figure 2A). With the other method based on density gradient centrifugation, we collected a total of 21 fractions and selected one “pure” fraction with almost no impurities (Figure 2B) and one “impure” fraction that contained mostly protein debris but very few vesicles overall (Figure 2C). The TEM images also show that isolated BMVs are very similar in size regardless of the isolation protocol used. The membrane-binding fluorescence dye FM4-64 has been used in the past to quantify bacteria-derived vesicles with a focus on the overall yield [13, 20]. After vesicle isolation, we analyzed whether there is a linear correlation between the vesicle amount and the fluorescence signal emitted by FM4-64. Therefore, we stained crude *P. aeruginosa* wild-type BMVs and measured the emitted signal of a 20-to 320-fold dilution series (Figure 3A). To compare BMV concentrations across different pools, we introduce the semi-quantitative unit “vesicle load” (VL). The goal was setting up a reference unit as a translation of the measured vesicle fluorescence signals for easier handling in dose-dependent cell culture experiments. The lowest vesicle dilution was set to a VL value of 1. The VL values of the remaining dilutions were set according to their dilution factor. We subtracted the fluorescence signal of control samples without vesicles and plotted the intensity difference in relation to the VL. The data clearly show a linear correlation between VL and fluorescence signal (*R*^2^ > 0.98), thus enabling the use of membrane staining dye for quantification of other BMV samples.

**Figure 2:**
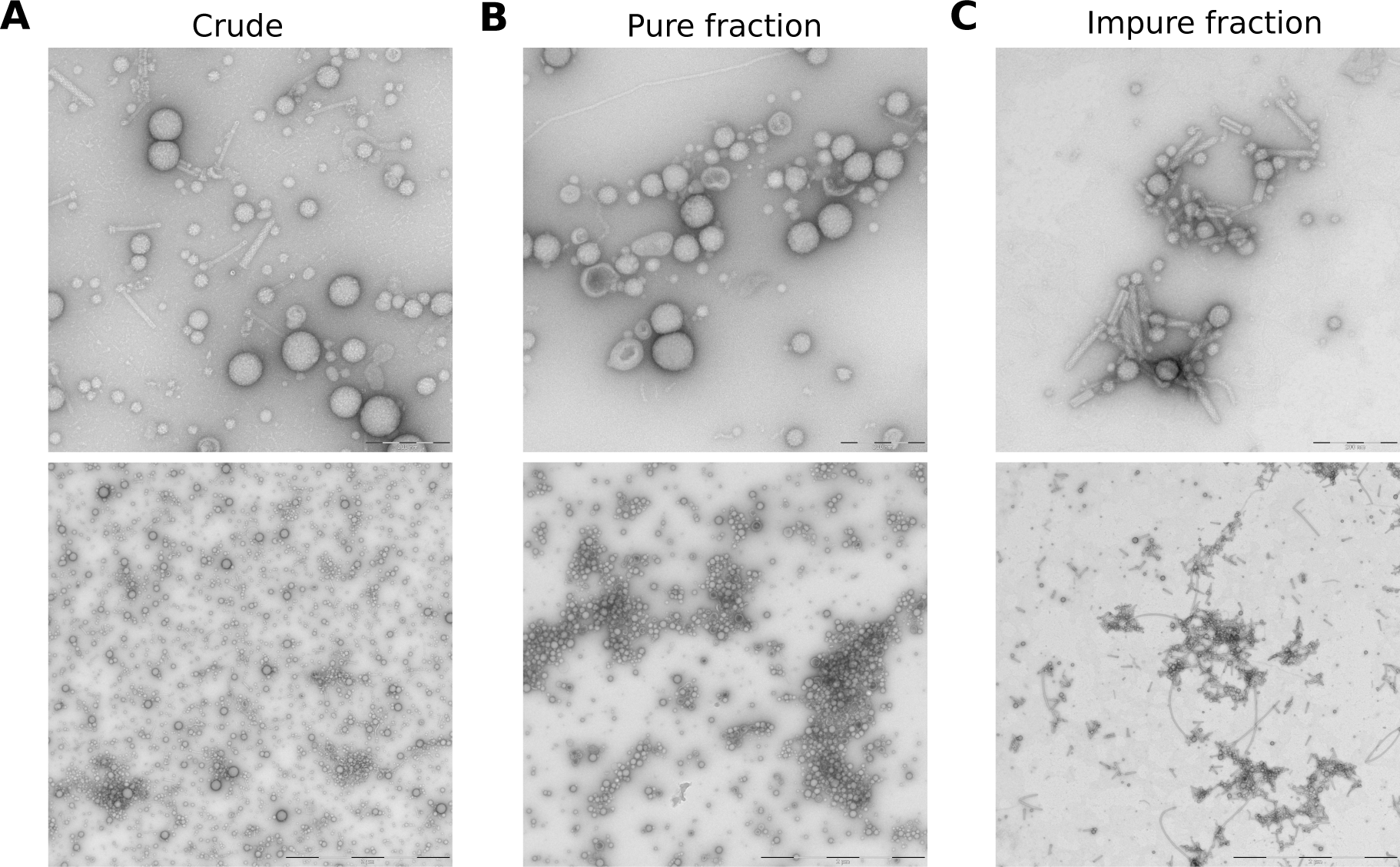
TEM images of different *P. aeruginosa* vesicle pools. **(A)** Crude wild-type BMVs were isolated using the fast protocol. **(B-C)** Wild-type BMVs were isolated with the iodixanol-based density gradient centrifugation protocol and stand representatively for a pure fraction without contaminants and an impure fraction containing mostly protein debris but very few vesicles. The majority of the visible impurities consists of pyocins and flagella parts. Scale bars = 200 nm (top), 2 µm (bottom).

**Figure 3:**
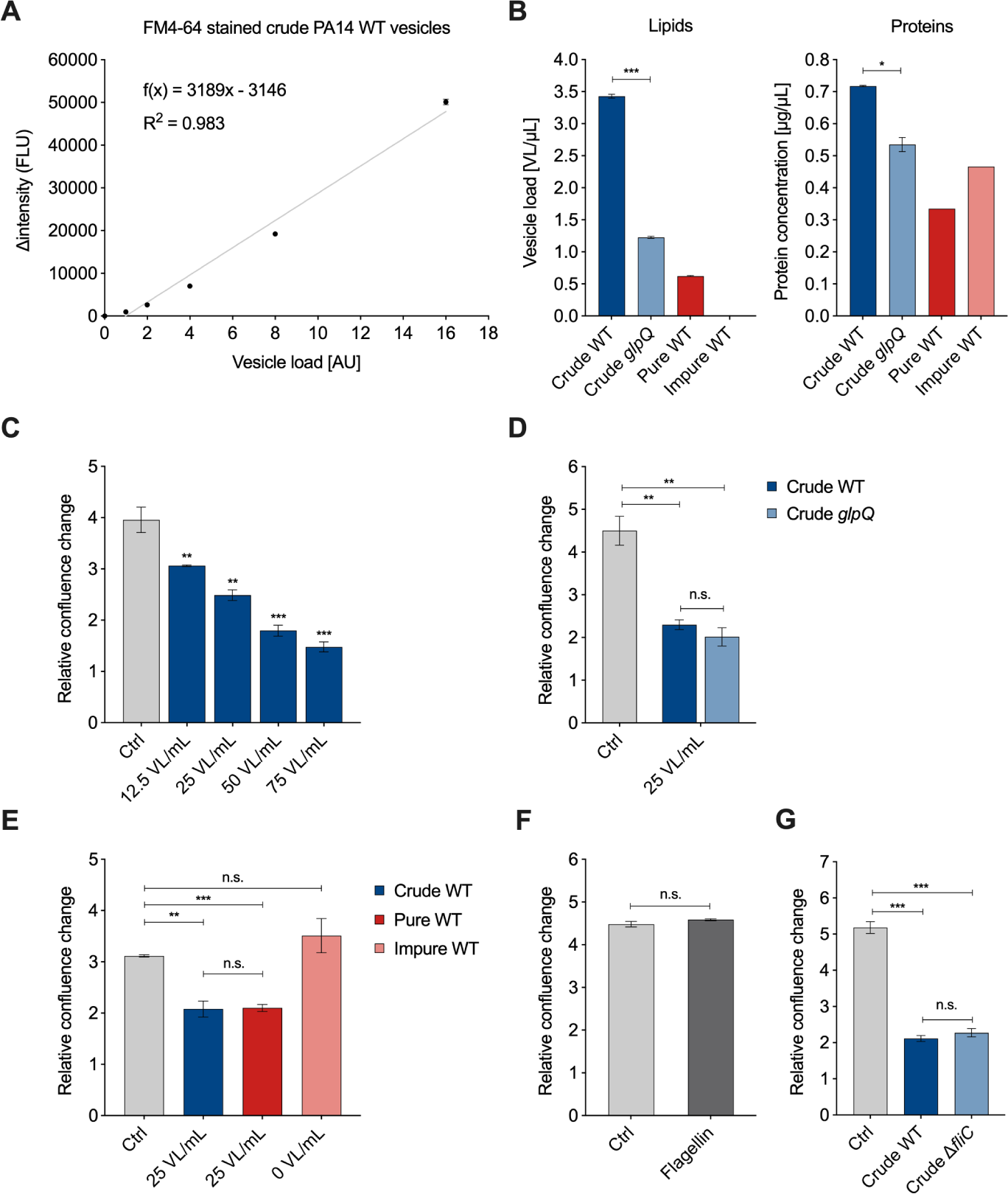
Quantification of *P. aeruginosa* BMVs and VL-dependent effects on A549 proliferation. **(A)** Linear regression of the fluorescence signal Δintensity (FLU) of FM4-64 stained crude *P. aeruginosa* wild-type vesicles in relation to the vesicle load (AU). Data were obtained from 3 technical replicates. **(B)** Lipid and protein concentrations of crude wild-type, crude *glpQ*, pure and impure wild-type BMV fractions. Lipid concentrations are defined as the determined vesicle load per volume (VL/µL) obtained from 3 technical replicates. Protein concentrations are obtained from 2 (*glpQ*) and 1 (fractions) technical replicates. **(C)** Relative confluence change of A549 cells treated with different concentrations of crude BMVs after 72 h. Data were obtained from 3 biological replicates. **(D)** Relative confluence change of A549 cells treated with crude wild-type and crude *glpQ* BMVs after 72 h. Data were obtained from 3 biological replicates. **(E)** Relative confluence change of A549 cells treated with crude, pure and impure wild-type BMVs after 72 h. Data were obtained from 3 biological replicates. **(F)** A549 proliferation after treatment with flagellin (1000 ng/mL) for 72 h. Data were obtained from 3 biological replicates. **(G)** Relative confluence change of A549 cells treated with crude wild-type and crude Δ*fliC* BMV after 72 h. Data were obtained from 3 biological replicates. All data in this figure are depicted as mean ± SEM. All significance niveaus were determined by Student’s t-test (n.s. = not significant, * = p < 0.05, ** = p < 0.01, *** = p < 0.001).

For the vesicle-host interaction experiments, we determined the VL and protein concentration of BMV pools from *P. aeruginosa* wild-type with varying purity and from the *glpQ* transposon mutant (Figure 3B). Compared to the crude wild-type BMVs, we observed that the VL concentration is lowered in the crude *glpQ* pool and pure wild-type gradient fraction. The fluorescence signal in the impure wild-type gradient fraction was below the limit of vesicle detection (blank level) resulting in a concentration of 0 VL/mL. These determined VL concentrations are in agreement with the results from TEM imaging (Figure 2). Moreover, we observed decreased protein concentrations in the crude *glpQ* transposon mutant as well as in both wild-type gradient fractions compared to the crude wild-type BMV pool (Figure 3B). We found a higher total protein concentration in the impure wild-type fraction than in the pure wild-type fraction, despite the latter having a higher VL. This underlines further that there is no direct correlation between protein and BMV concentration.

To demonstrate the benefit of normalizing the vesicle amount by the VL in vesicle-host interaction experiments, we analyzed the effect of different VL concentrations on the proliferation of A549 lung cancer cells. It is well known that BMVs of many pathogenic and non-pathogenic bacteria reduce cell proliferation of cancer cells *in vitro* and in *in vivo* tumor models [21, 22]. In line with these results, we observed that the proliferation of A549 cells was decreased in a VL-dependent manner (Figure 3C). For all following experiments, we decided to use a concentration of 25 VL/mL, as this already significantly reduce cell proliferation. Next, we examined whether crude wild-type vesicles and crude vesicles derived from the *glpQ* transposon mutant affected cell proliferation similarly. Indeed, vesicles of both strains reduced cell proliferation to the same extent when normalized to 25 VL/mL. Interestingly, the VL of the crude *glpQ* mutant-derived vesicles was around three times lower compared to the wild-type vesicles, whereas the protein concentration was only slightly reduced (Figure 3B), which equals more overall protein delivery in the VL-normalized experiments. This result clearly shows that the effect of these vesicles on cell proliferation is independent of the total protein concentration. Moreover, it demonstrates that GlpQ itself does not contribute to the vesicle-induced proliferation reduction, but it might affect overall vesicle generation and protein incorporation by *P. aeruginosa*.

As described above, crude vesicles contained common impurities like flagella and pyocins. For this reason, we next analyzed how much these contaminants contribute to the observed cell proliferation effect and repeated the proliferation experiment with the aforementioned pure and impure wild-type gradient fractions. Because the VL was below the detection limit in the impure wild-type fraction, it was not possible to normalize it to 25 VL/mL. Instead, we used the volume corresponding to the pure wild-type fraction. Normalized amounts of crude and pure wild-type vesicles reduced A549 proliferation similarly, whereas the impure wild-type fraction did not reduce cell proliferation significantly (Figure 3E), despite having a higher overall protein content compared to the pure wild-type fraction. This observation matches the low vesicle count shown in the TEM images as well as the VL being below the detection limit, further underlining the better performance of VL over total protein amounts. After showing that the impure wild-type fraction, even though containing impurities, had no significant impact on A549 proliferation, we analyzed whether we would observe similar results with purified flagellin protein from *P. aeruginosa*. Flagellin protein is the main structural component of flagella and essential for their formation. A549 cells treated with a high dose of flagellin (1000 ng/ml) showed no significant difference from the control (Figure 3F). To further investigate the effect of flagellin, we used the knockout mutant strain Δ*fliC* which does not form flagella due to the lack of the filament protein and isolated crude BMVs with the fast protocol. Again, we did not observe a significant difference compared to the crude wild-type vesicles (Figure 3G). Taken together our results suggest that contaminants like flagella, despite their known immunogenic potential [23, 24], do not significantly affect the proliferation of lung epithelial cancer cells.

After showing that the VL gives reproducible results for cell proliferation experiments, we next thought to analyze whether the same is true for the vesicle-induced effects on cellular metabolism. To this end, we performed stable isotope-assisted metabolomics using [U-^13^C_6_]glucose and determined intracellular metabolite levels as well as the isotopic enrichment in the form of mass isotopomer distributions (MIDs). We treated A549 cells with either *P. aeruginosa* wild-type vesicles (crude and pure) and crude *glpQ* mutant vesicles using 25 VL/mL. First, we analyzed the intracellular metabolite levels, mainly citric acid cycle intermediates and amino acids (Figure 4A). Overall, we observed that all vesicle-treated samples clustered together regardless of their purity or strain origin. We found that all citric acid cycle intermediates and those closely connected to it were less abundant in vesicle-treated A549 cells. On the other hand, we observed that almost all amino acid levels were increased, most predominantly in cells treated with pure vesicles. Several studies have shown that bacterial-derived vesicles are associated with the induction of autophagy and mitophagy in host cells [10, 25–27] which might lead to the observed increase of amino acid levels.

**Figure 4:**
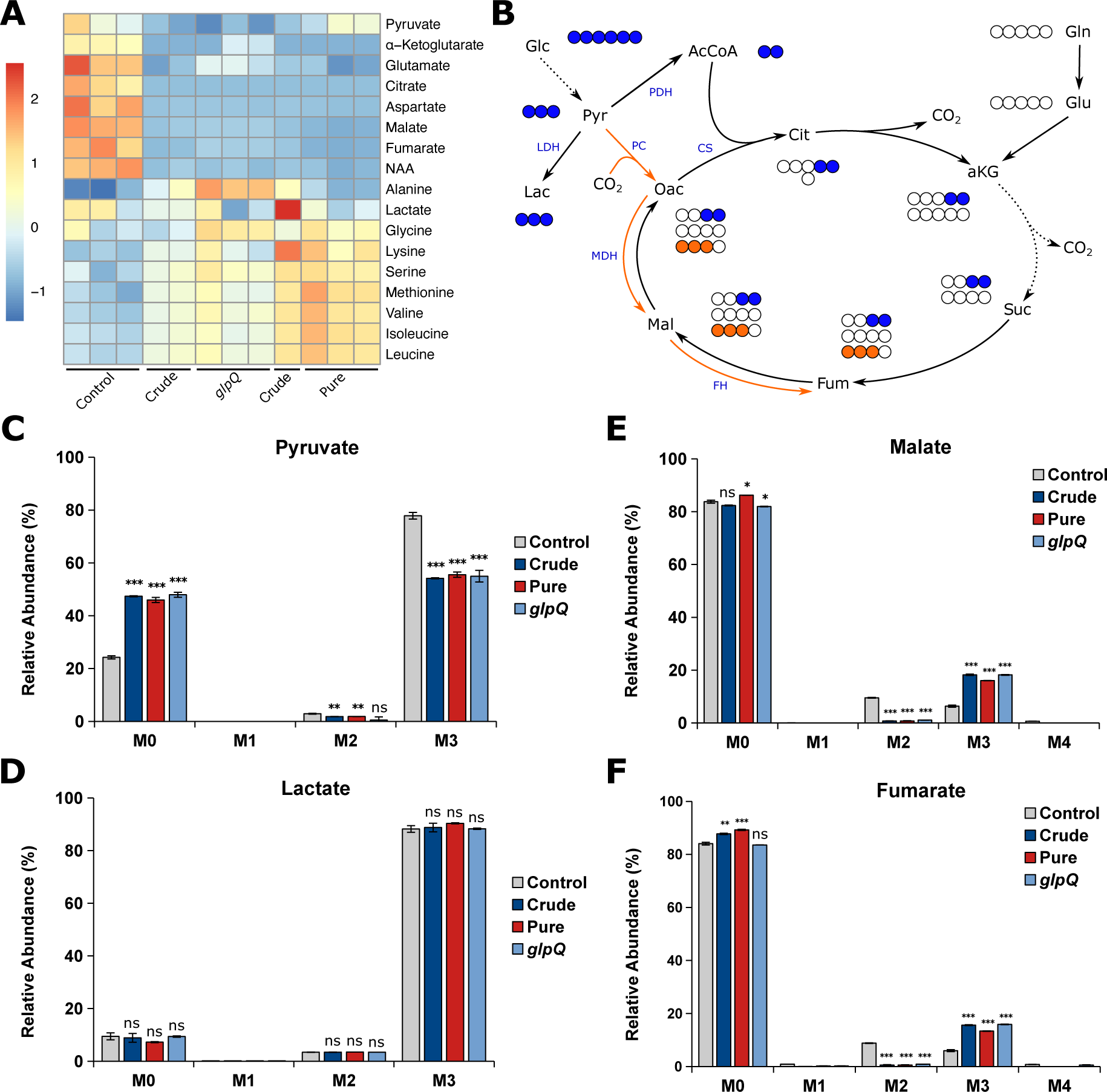
*P. aeruginosa* vesicles alter the metabolism of A549 cells. **(A)** Heatmap of intracellular metabolite levels of A549 cells treated with different vesicle pools measured with GC-MS. Metabolite levels were normalized to cell confluence and z-score standardized. **(B)** Schematic overview of carbon atom transitions from [U-^13^C_6_]glucose. Glc, glucose; Pyr, pyruvate; Lac, lactate; AcCoA, acetyl coenzyme A; Cit, citrate; aKG, *α*-ketoglutarate; Suc, succinate; Fum, fumarate; Mal, malate; Oac, oxaloacetate; Gln, glutamine; Glu, glutamate; LDH, lactate dehydrogenase; PDH, pyruvate dehydrogenase; PC, pyruvate carboxylase; CS, citrate synthase; MDH, malate dehydrogenase; FH, fumarate hydratase. **(C-F)** Relative mass isotopomer abundances of pyruvate, lactate, fumarate, and malate. M0 - M4 represent the amount of labeled carbon atoms derived from [U-^13^C_6_]glucose in a single molecule. A549 cells were incubated for 24 h. Data were obtained from 3 biological replicates and are depicted as mean ± SEM. All significance niveaus were determined by Student’s t-test (n.s. = not significant, * = p < 0.05, ** = p < 0.01, *** = p < 0.001).

In addition to the intracellular metabolite levels, we analyzed vesicle-induced intracellular flux changes and whether the VL is suitable to create reproducible results. Usually, the majority of pyruvate is derived from glycolysis, leading to M3 isotopologues from [U-^13^C_6_]glucose (Figure 4B). We observed decreased M3 pyruvate for all vesicle-treated cells, suggesting that pyruvate is additionally derived from another carbon source other than glucose (Figure 4C). Interestingly, the labeling pattern of lactate did not change upon vesicle treatment (Figure 4D), suggesting that the increase of unlabeled pyruvate is not derived from glycolysis. Because we observed reduced levels of TCA cycle-related metabolites, we expected a reduced oxidative TCA cycle flux. Glucose carbons can enter the TCA cycle either via pyruvate dehydrogenase complex (PDH) or pyruvate carboxylase (PC) activity, leading to distinct labeling patterns (Figure 4B). Briefly, PDH activity generates [^13^C_2_]acetyl-CoA from which M2 malate is derived via the oxidative TCA cycle, whereas PC activity yields [^13^C_3_]oxaloacetate leading to M3 malate catalyzed by malate dehydrogenase (MDH). We found that M2 malate and fumarate significantly decreased to almost zero in vesicle-treated cells. On the other hand, we observed a significant increase in M3 malate and fumarate in BMV-treated cells as a direct result of higher PC activity (Figures 4E and 4F). Taken together, we observe a reduced oxidative TCA cycle flux upon vesicle treatment, which probably leads to the aforementioned decrease in TCA cycle-related metabolite levels. These alterations in mitochondrial metabolism are not surprising, as BMV-induced mitochondrial dysfunction is already known for macrophages, where BMVs activate intrinsic mitochondrial apoptosis and inflammation [28].

For all of the aforementioned metabolic flux changes, we observed little to no difference whether BMVs were isolated from a *P. aeruginosa* wild-type or *glpQ* transposon mutant strain. Additionally, we observed the same effect for crude and pure wild-type BMVs, meaning that the method of isolation and thus the purity of the vesicle sample did not have an significant impact on vesicle-induced metabolic changes in A549 cells.

Finally, we intended to show that the VL can be used to compare vesicles from different strains, as proposed in Figure 1. To that end, we selected five of the most abundant *P. aeruginosa* vesicle proteins and isolated crude BMVs of the corresponding transposon mutant strains: The aforementioned glycerophosphoryl diester phosphodiesterase (*glpQ*), inositol-1-monophosphatase (*suhB*), aminopeptidase (*lap*), lactonizing lipase (*lipA*) and structural outer membrane porin precursor (*oprF*). Using a concentration of 25 VL/mL, all mutant strain-derived vesicles reduced cell proliferation of A549 cells, except vesicles lacking SuhB (Figure 5), suggesting that it is contributing to the observed proliferation effects of *P. aeruginosa* BMVs. Our results are in line with previous studies that have already shown an essential role of *suhB* for regulating multiple virulence genes and motility of *P. aeruginosa* [29, 30], but so far it had never been associated with vesicle biogenesis and their functionality in terms of virulence. In conclusion, this experiment demonstrates the possibility of VL normalization for the analysis of BMVs isolated from different mutant strains.

**Figure 5:**
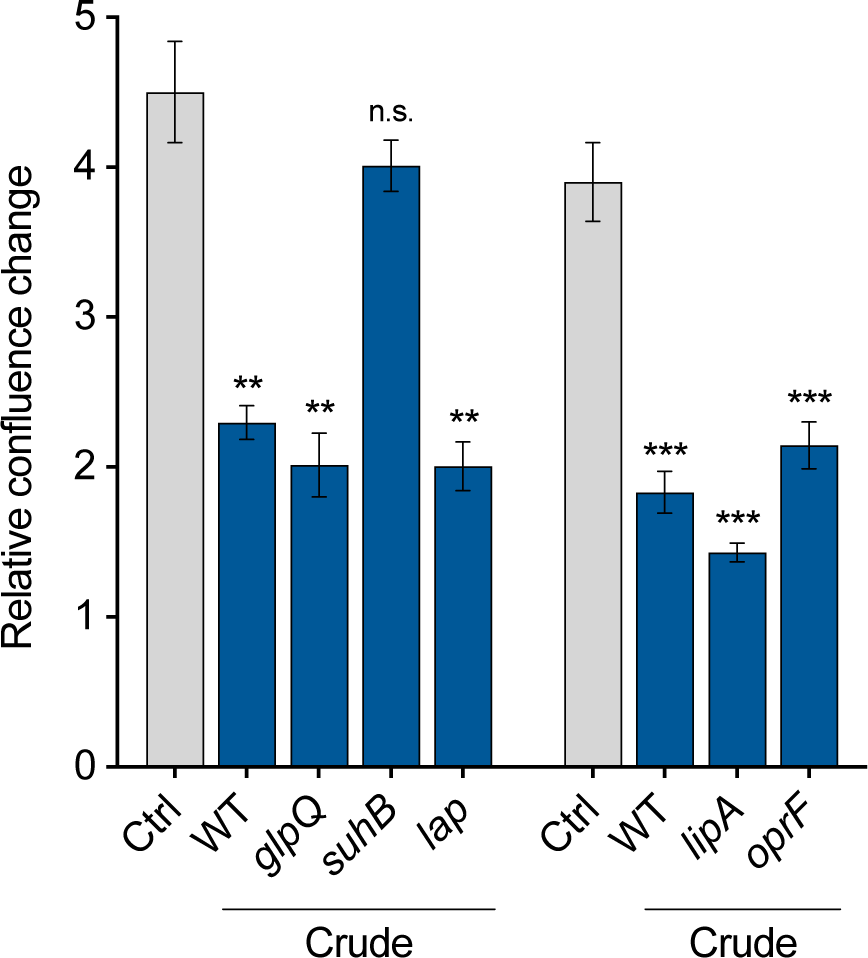
A549 proliferation after treatment with vesicles isolated from *P. aeruginosa* transposon mutants. Proliferation of A549 cells treated with crude vesicles of different *P. aeruginosa* transposon mutants lacking high abundant BMV-associated proteins (25 VL/mL, 72 h). Data were obtained from 3 biological replicates and depicted as mean ± SEM. All significance niveaus were determined by Student’s t-test (n.s. = not significant, * = p < 0.05, ** = p < 0.01, *** = p < 0.001).

In summary, we have shown that the VL works very well to perform reproducible vesicle cell culture interaction experiments. However, it is worth noting that the VL is a semi-quantitative quantification method as it depends on the vesicle concentration of the sample used for the initial regression analysis. It does not provide absolute quantification which would require a vesicle standard with a known concentration. Though, semi-quantification is sufficient for most applications. We demonstrated several applications for the VL in vesicle-host interaction experiments with mammalian cell culture. We showed that the VL is reliable for the quantification of different *P. aeruginosa* BMV pools including transposon mutants and that BMV treatment of A549 cells with VL normalization leads to reproducible proliferation reductions and metabolic flux changes. Moreover, our results elucidated that the protein SuhB is an important contributor of vesicle-induced pathogenesis.

## SUPPLEMENTARY INFORMATION

**Figure S1:**
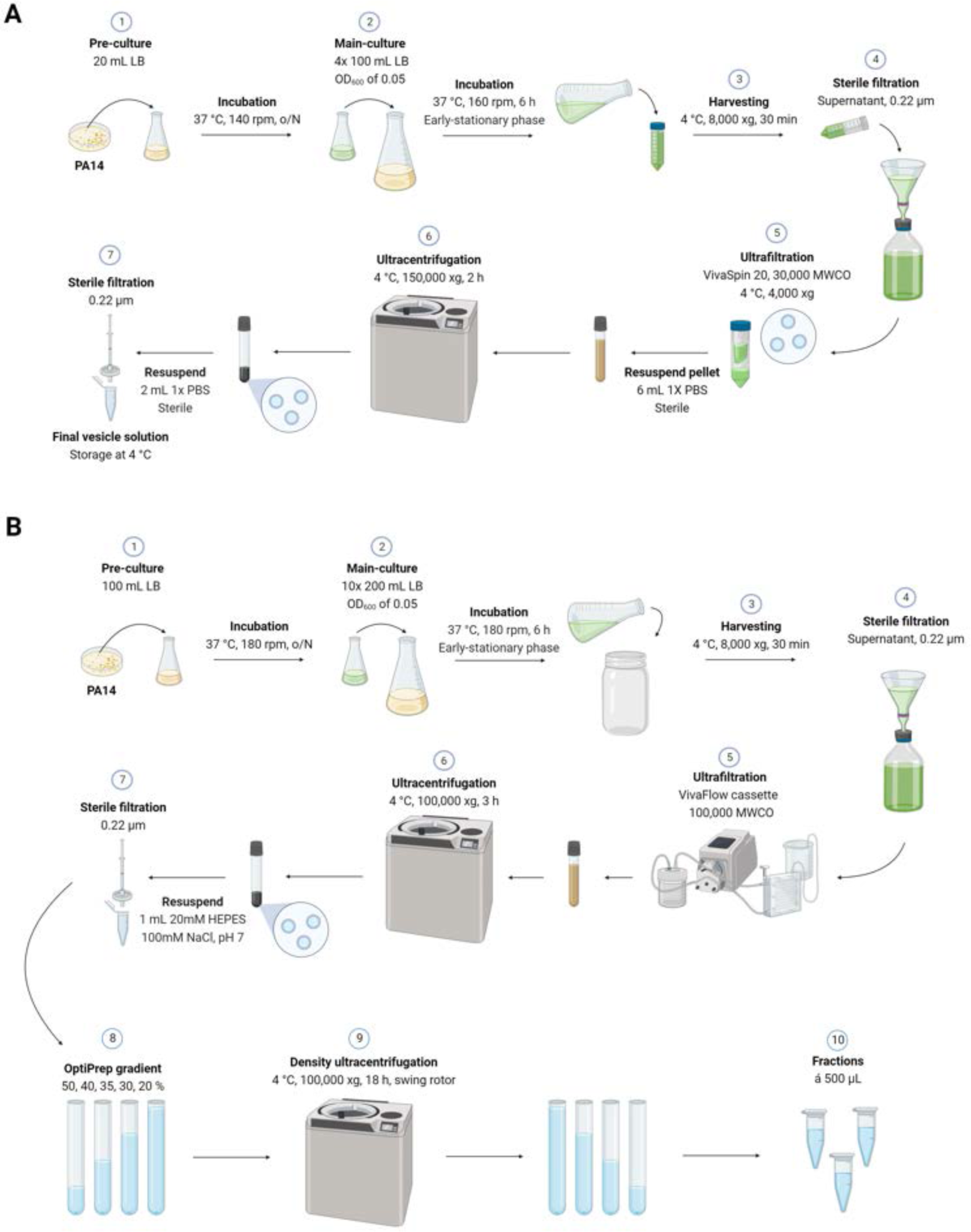
Isolation workflow of crude and pure *P. aeruginosa* BMVs. (**A**) Crude isolation of BMVs by ultrafiltration combined with ultracentrifugation: (1) Bacterial pre-cultures are grown overnight in LB broth (+ needed supplements like antibiotics). (2) Main cultures are inoculated with an OD600 of 0.05 and incubated until they reach the early-stationary phase of growth. (3) Bacterial cells are harvested and the culture supernatant filtered. (5) Ultrafiltration of sterile culture supernatants leads to concentrated material. (6) Membrane vesicles are pelleted by ultracentrifugation of concentrates and resuspended in a buffer. (7) Membrane vesicle solutions are sterile filtered and stored at 4 °C. **(B)** Pure isolation of BMVs based on density gradient centrifugation. The first steps (1-7) are performed similar to the crude isolation method. Afterwards, an iodixanol density gradient is used to remove remaining contaminants and get the final vesicle fractions (8-10). Created with biorender.com.

**Figure S2:**
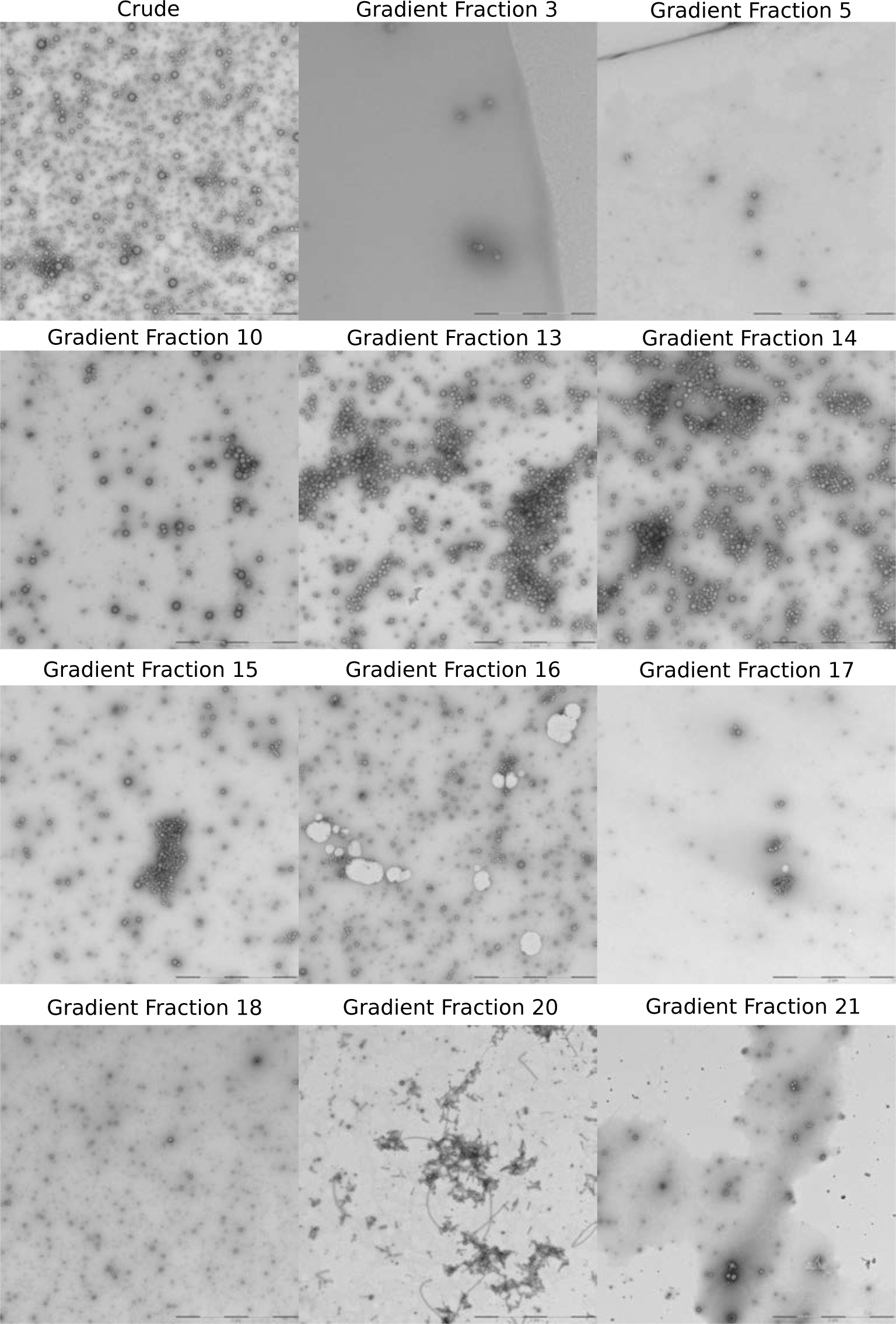
TEM images of selected negative stained *P. aeruginosa* BMVs. TEM images of negative stained crude and pure vesicles. Crude wild-type BMVs were isolated using the fast protocol. The wild-type vesicle fractions were isolated with the iodixanol-based density gradient centrifugation protocol. Scale bars = 2 µm.

